# Polymicrobial extracellular vesicles reduce the innate immune response of human cystic fibrosis bronchial epithelial cells

**DOI:** 10.64898/2026.04.09.717493

**Authors:** Lily A. Charpentier, Roxanna L. Barnaby, Carolyn T. Roche, Byoung-Kyu Cho, Prashant Kaushal, Young Ah Goo, Brad Vietje, Douglas J. Taatjes, Alix Ashare, Fabrice Jean-Pierre, Bruce A. Stanton

## Abstract

Chronic antibiotic-resistant cystic fibrosis (CF) lung infections are the leading cause of death in adults with CF. Despite advances in highly effective modulator therapies, microbial communities persist in the CF lung. The pathogenesis of CF airway infections can be exacerbated by pathogens such as *Pseudomonas aeruginosa,* which communicates with primary human bronchial epithelial cells (pHBEC) by secreting bacterial extracellular vesicles (bEVs) that diffuse through mucus and deliver virulence factors, DNA, and RNA to pHBEC. However, most CF lung infections are polymicrobial in nature, and therefore, the contribution of polymicrobial bEVs remains to be determined. By using a polymicrobial culture model representing a ‘pulmotype’ detected in ∼34% of lung infections in people with CF (pwCF), comprised of *P. aeruginosa*, *Staphylococcus aureus*, *Streptococcus sanguinis*, and *Prevotella melaninogenica* grown in synthetic sputum medium under anoxia, we report that each bacterial genus in the polymicrobial community secretes bEVs containing proteins and RNAs predicted to promote the establishment of chronic infection by enhancing virulence, biofilm formation, and upregulating the stress response and pro-inflammatory pathways in pHBEC. This response is most pronounced in CF pHBEC. Elexacaftor/Tezacaftor/Ivacaftor (ETI), a highly effective modulator therapy, does not ameliorate the response or return it to WT levels. Bacterial EVs also inhibited ETI CFTR Cl^-^ currents by CF pHBEC. These studies provide insight into why ETI does not eliminate polymicrobial lung infections and a hyperinflammatory lung environment in pwCF.

**IMPORTANCE:** Cystic fibrosis (CF) is a genetic disease characterized by chronic polymicrobial lung infections that, if untreated, are one of the primary causes of death in CF. Elexacaftor/Tezacaftor/Ivacaftor (ETI) has many positive clinical outcomes, but it does not eliminate chronic polymicrobial lung infections or inflammation. Using a new biologically relevant co-culture model, we have demonstrated that bacteria secrete vesicles (bEVs) that contain proteins and RNAs. We observed that these RNA-loaded bEVs are predicted to promote the pathogenesis of chronic CF lung infections by enhancing bacterial virulence and biofilm formation, as well as upregulating the pro-inflammatory response in lung cells. ETI does not ameliorate the response of lung cells to bEVs. Our research will facilitate the development of more effective approaches to eliminate infection and inflammation in CF and other lung diseases characterized by chronic polymicrobial infections and excessive inflammation.

## INTRODUCTION

Cystic fibrosis (CF) is a disease that affects over 100,000 people worldwide (1) and is caused by mutations in the cystic fibrosis transmembrane conductance regulator gene (CFTR), which encodes a chloride and bicarbonate ion channel. It is a multiorgan disease characterized by the formation of thick hypoxic/anoxic mucus in the lungs, leading to low mucociliary clearance, the establishment of chronic, antibiotic-tolerant lung infections, and excessive inflammation (2–4). In people with CF (pwCF), lungs are initially colonized by *Staphylococcus aureus*, which can be cleared by antibiotics, but later evolves into chronic, antibiotic-tolerant polymicrobial infections (3–11). Highly effective modular therapies (HEMT) have been developed to improve CFTR-mediated chloride and bicarbonate secretion by airway epithelial cells. One of the most widely used HEMTs, Elexacaftor/Tezacaftor/Ivacaftor (ETI) (12), reduces inflammation and stimulates antimicrobial peptide secretion by CF primary human bronchial epithelial cells (pHBEC) (13,14). Although ETI is clinically beneficial and improves lung function, it does not eliminate chronic polymicrobial infections or the hyperinflammatory lung environment (2,3,5,15–21). Thus, new models are needed to elucidate the effects of polymicrobial infections on CF pHBEC and to develop new approaches to eliminate lung infections and inflammation in pwCF.

Although *Pseudomonas aeruginosa* is considered a canonical CF pathogen and chronically infects ∼35% of adults with CF, the overwhelming majority have polymicrobial lung infections (22,23). The utilization of a recently developed CF microbiome-informed polymicrobial community model −representing ∼34% of lung infections in pwCF− composed of *P. aeruginosa*, *S. aureus*, *Streptococcus* and *Prevotella* spp. grown in synthetic CF medium (SCFM2) under anoxic conditions has shown that interspecies interactions can impact CF-relevant bacterial phenotypes such as antibiotic sensitivity (22,24–27). Several studies have reported the co-existence of multiple bacterial species in the CF lung, and their co-detection is correlated with a negative impact on patient outcomes (23,28). More importantly, while single variables, including one bacterial genus, Simpson diversity, and age, explain less than 11% of patient variability in lung function, the presence of polymicrobial communities accounts for 27% of this variability (22,23). *P. melaninogenica*, an anaerobe, is included as anaerobes are important contributors to positive CF outcomes (29–32). Moreover, this selection of bacteria mimics the metabolite cross-feeding relationships between these four genera (28). So far, this validated community model has mainly been used to probe questions focused on elucidating mechanisms of interaction among its members. A few studies have focused on host-pathogen interactions in coculture with two of the four species. For example, studies of *P. aeruginosa* and *S. aureus* have found that coculture increases both antibiotic resistance and bacterial internalization into lung epithelial cells (33). *S. aureus* also reduces the *P. aeruginosa*-induced host cell IL-8 response (34). *Streptococcus sanguinis* uses reactive nitrogen species to antagonize *Prevotella melaninogenica* within the CF respiratory microbiome (26). However, to the best of our knowledge, no studies have reported how the presence of CF-relevant polymicrobial communities containing all four bacteria affects the biology of wild-type (WT) and CF pHBEC.

Most bacteria in the lungs reside in hypoxic/anoxic CF mucus plugs that overlie lung epithelial cells (35). A steep oxygen gradient forms in CF mucus, with oxygen permeating about 1 μm into the sputum, which is ∼20 μm thick, before becoming anoxic (35–38). In the model used for this study, bacteria are grown in an anoxic environment in artificial CF sputum (SCFM2), and include an obligate anaerobe, *P. melaninogenica*, to reflect the CF lung environment (22). However, as anoxia damages bronchial epithelial cells, our study examines the effect of bacterial extracellular vesicles (bEVs) secreted by this polymicrobial community on pHBEC grown in the presence of oxygen, as described (39). This approach enables a study of bEVs on pHBEC without causing low-oxygen stress. Our study fills a significant knowledge gap and provides an opportunity to identify potential therapeutic targets in host-microbe interactions in biologically relevant conditions.

We have shown previously that *P. aeruginosa* communicates, in part, with lung epithelial cells by secreting bEVs that diffuse through mucus and fuse with lipid rafts on the lung epithelial cell surface. These vesicles deliver virulence factors, small RNAs (sRNAs), and tRNA fragments to pHBEC that reduce CFTR Cl^-^ secretion (40–42). Furthermore, bEVs suppress the inflammatory response to lipopolysaccharides (LPS), thereby disrupting immune cell recruitment to sites of infection and bacterial clearance (43). Taken together, the aim of this study is to begin elucidating how bEVs secreted by a representative CF polymicrobial community affect the innate immune response of WT and CF pHBEC, as well as the cellular responses of CF pHBEC to ETI.

We demonstrate that each bacterial species in this CF polymicrobial community secretes bEVs containing proteins, sRNA, and tRNA fragments predicted to promote the establishment of chronic infection by enhancing bacterial virulence and biofilm formation, and by upregulating the stress response and inflammatory pathways in pHBEC. This response is most pronounced in CF pHBEC. Interestingly, ETI does not return the CF pHBEC response to bEVs to WT levels, highlighting the need for continued research into novel therapeutics to reduce chronic, antibiotic-resistant polymicrobial infections.

## RESULTS

### bEVs are secreted in a CF-relevant polymicrobial community

bEVs secreted by the polymicrobial culture containing *P. aeruginosa*, *S. aureus*, *S. sanguinis*, and *P. melaninogenica* were isolated using OptiPrep density-gradient ultracentrifugation, as recommended by the International Society for Extracellular Vesicles (44). Nine resulting fractions were examined for protein content (**Fig. 1A**). As Fractions 1 and 2 displayed the highest protein concentrations, these fractions were pooled for downstream analyses of bEVs. The bEVs contain 2.8×10^14^ particles/µg protein. More than 3×10^9^ particles/µg of protein is considered a standard of purity for bEV preparations (45). High-resolution negative-stain transmission electron microscopy (TEM) was employed to visualize bEVs from the pooled fractions alongside a processed control (PC; uninoculated media that had undergone identical density-gradient isolation) (**Fig. 1B–H**). Vesicle diameters determined by TEM ranged from ∼20 nm to ∼250 nm, with the majority falling within 40–90 nm (**Fig. 1B**), consistent with previously reported bEV sizes for each species (44,46,47). This size distribution is smaller than measurements obtained by nanoparticle tracking analysis (NTA) (167.6 ± 23.8 nm, *N* = 5 independent experiments). The discrepancy is likely due to the NTA’s tendency to overestimate diameter due to measuring light scattering (48), while the sample preparation for TEM causes the shrinkage of vesicles (49). Heterogeneous vesicular structures were evident in the pooled fractions (**Fig. 1E–H**) but were absent in the processed control (**Fig. 1C–D**). Branched structures, pictured in **Fig. 1C-D**, reminiscent of mucin from the SCFM2, were very occasionally observed. Moreover, the vesicles in **Fig. 1D** displayed a similar morphology to that we previously observed for *P. aeruginosa* vesicles in monoculture (39).

**Fig. 1.**
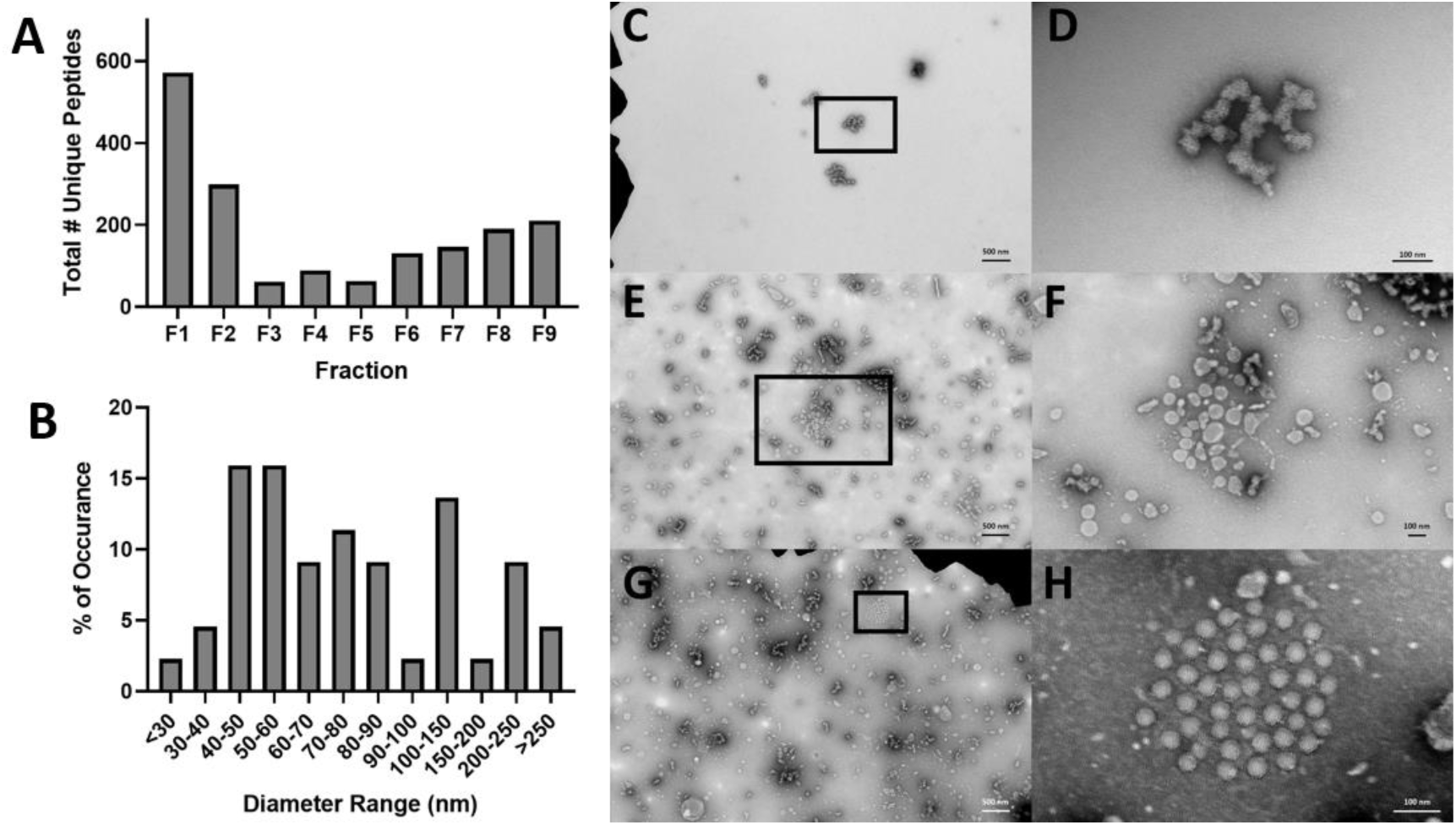
Isolation and characterization of bEVs. A) Peptide counts in each of 9 fractions of the OptiPrep gradients. B) Size distribution analysis of bEVs in fractions F1 and F2 as determined by transmission electron microscopy. C) SCFM2, processed control (magnification 6,000x). The area contained within the box in C is presented at higher magnification in (D) (40,000x). E) Polymicrobial bEVs (magnification 6,000x). F) Polymicrobial bEVs (magnification 20,000x). The area contained within the box in E is presented at higher magnification in F. (G) Polymicrobial bEVs with a structure reminiscent of *P. aeruginosa* outer membrane vesicles (magnification 6000x) (8). The area contained within the box G is presented at higher magnification in H (magnification 50,000x).

In accordance with recommendations by the ISEV (44), the polymicrobial bEVs were evaluated using multiple orthogonal methods. We have previously demonstrated that bEVs contain 16S rRNA (43). Parallel 16S rRNA sequencing and relative fluorescence measurements of the bEV preparation confirmed the presence of bEVs secreted by all four bacterial species (**Fig. 2**). Although the bacteria grew to comparable colony-forming unit (CFU) densities (**Fig. 2A**), the concentration of bEVs differed among species (**Fig. 2B**). Across both sequencing-based and fluorescence-based approaches, *P. aeruginosa* secreted the highest number of bEVs, followed by *S. sanguinis*, *S. aureus*, and *P. melaninogenica* (**Fig. 2B**). To validate the clinical relevance of the polymicrobial bEV model, bEVs were isolated from the bronchial alveolar lavage fluid (BALF) of two pwCF (*Phe508del/Phe508del*, post-ETI) and subjected to 16S sequencing to quantify the average relative abundances of vesicle-associated bacterial genera in the donors’ BALF. All four species comprising our *in vitro* model were identified among the ten most relevant bEV genera in donor BALF (**Fig. 2C**). Together, they comprise 53.3% of the bEVs represented in the BALF (**Fig. 2C**).

**Fig. 2.**
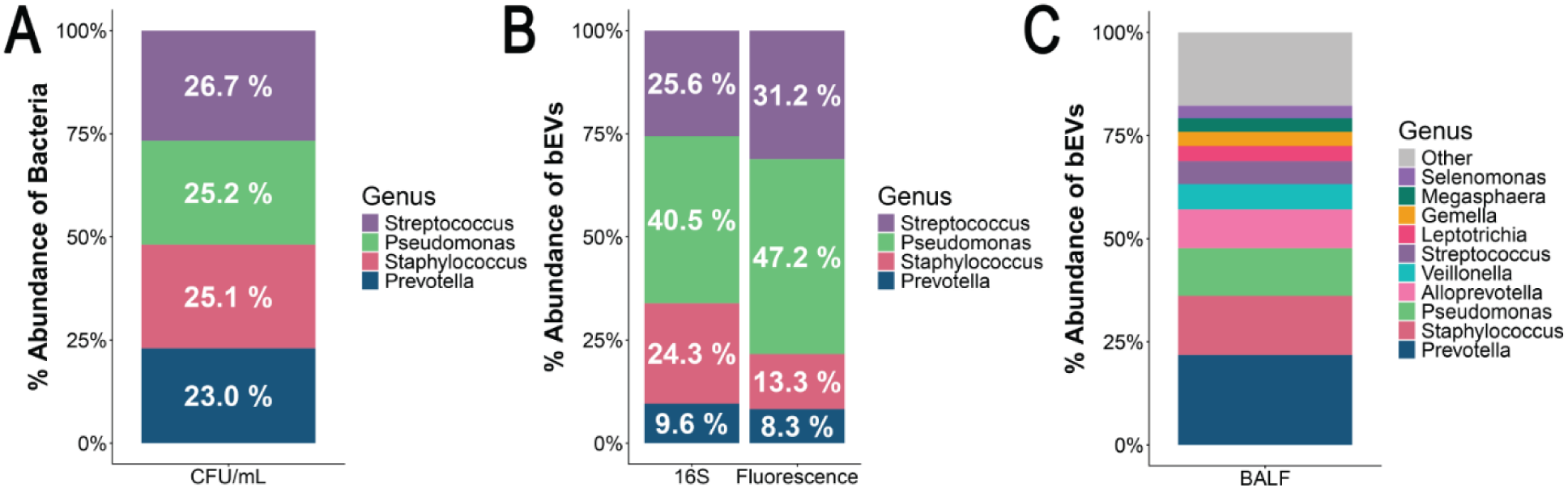
All four species present in the polymicrobial culture release bEVs. A) Proportion of each species of bacteria in culture as determined by CFU/mL at the time of bEV isolation. *N*=3 independent experiments. B) Proportion of bEVs present in the polymicrobial culture as determined by 16S rRNA sequencing and fluorescence of bEV preparations measured using Zetaview NTA. *N*=3. C) Proportions of top genera in CF BALF by 16S sequencing. bEVs secreted from all four genera in the polymicrobial culture were also identified in clinical samples obtained from pwCF. CF BALF was obtained from banked samples from the Dartmouth CF RDP Translational Research Core with approval from the Dartmouth Health IRB.

### Polymicrobial bEVs contain proteins that facilitate biofilm formation, inter-bacterial signaling, and enhanced virulence

To identify the biological processes and pathways represented by the proteins packaged within the polymicrobial bEVs, we performed a pathway-activation analysis using ESKAPE Act Plus (50). A total of 41 significantly enriched KEGG pathways (*P*<0.05) in *P. aeruginosa* bEVs, 16 in *S. aureus* bEVs, 31 in *S. sanguinis* bEVs, and the Ribosome pathway in *P. melaninogenica* were identified (**Fig. 3, Table S1**). Across all enrichment analyses, several known virulence-associated pathways emerged as consistently significant. The biofilm-formation and bacterial chemotaxis pathways were prominent, with significant enrichment observed in *P. aeruginosa* (**Fig. 3A**). The quorum-sensing pathway was also enriched in *P. aeruginosa*, *S. aureus*, and *S. sanguinis* bEVs, highlighting the potential of bEVs to influence community-wide communication and formation of antibiotic-resistant biofilms (**Fig. 3A-C**). Additionally, the β-lactam resistance pathway was enhanced in *P. aeruginosa* and *S. sanguinis* (**Fig. 3A** and **3B**). This analysis indicates that each bacterial species secretes proteins in bEVs that are involved in pathways associated with infection and antibiotic resistance (**Fig. 3, Table S1**).

**Fig. 3.**
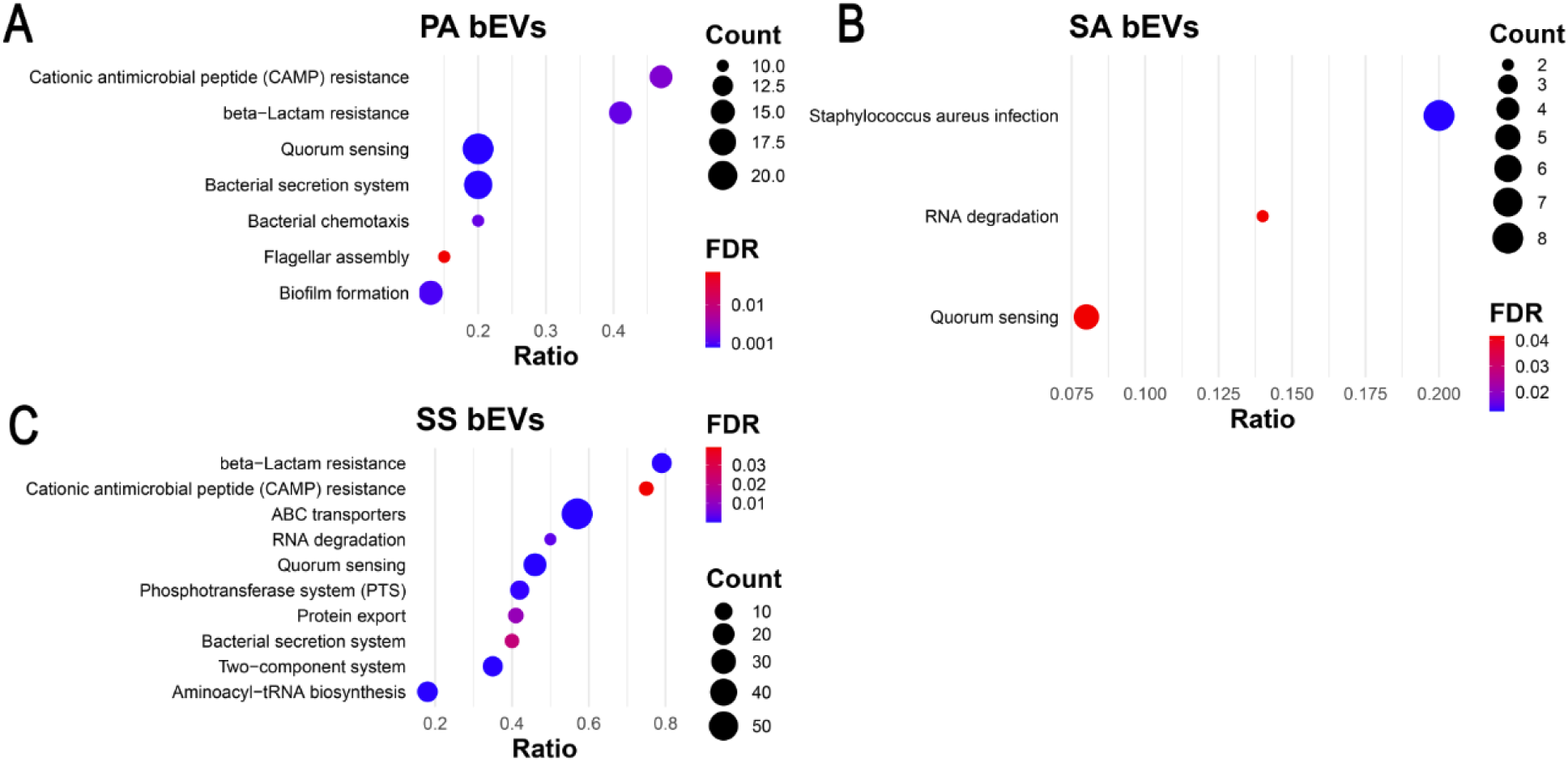
Polymicrobial bEVs contain proteins associated with CF lung infection pathogenesis. KEGG Pathway enrichment analysis of antibiotic resistance and virulence pathways for proteomic data from bEVs secreted by the polymicrobial culture. A) *P. aeruginosa* (PA). B) *S. aureus* (SA). C) *S. sanguinis* (SS). KEGG pathway analysis for *P. melaninogenica* could not be performed due to a lack of available pathway data. Count is the number of proteins identified as part of the pathway. FDR, False Discovery Rate. Ratio is the number of identified proteins divided by the total number of proteins in the KEGG pathway.

Proteomic analysis of bacteria in the polymicrobial community was also conducted to compare KEGG pathway analysis of bacteria versus bEVs. Analysis of the data revealed a similar list of protein networks in bacteria and bEVs. In *P. aeruginosa*, the 41 significant KEGG pathways observed in bEVs were also found in the bacteria (**Fig. 4, Table S2**). In *S. aureus,* KEGG pathways identified in bEVs were also widely represented, with only the RNA degradation pathway not enhanced in *S. aureus* bacteria, suggesting selective packaging of proteins in bEVs, which has been described in previous studies (51,52). Similarly, 30 of 31 bEV KEGG pathways were represented in *S. sanguinis*, with only the cationic antimicrobial peptide resistance pathway not identified in the bacteria. The small differences in KEGG pathways between bacteria and bEVs are most likely due to larger variances in protein abundance in bacteria compared to the bEVs. Collectively, KEGG pathway analysis revealed that bacteria and their bEVs share pathways that facilitate biofilm formation, inter-bacterial signaling, and enhanced virulence.

**Fig. 4.**
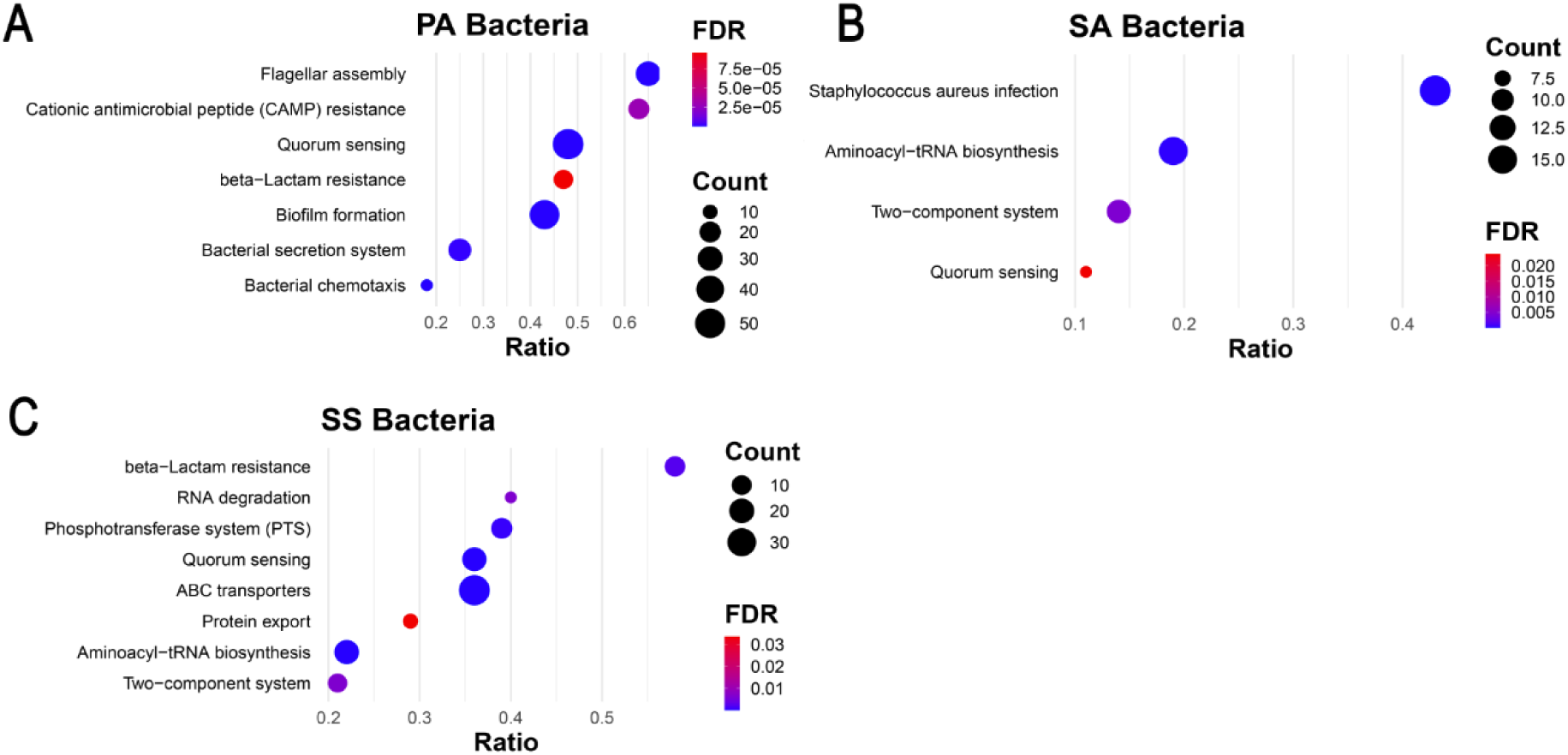
The polymicrobial culture contains proteins that promote infection. KEGG Pathway enrichment analysis of antibiotic resistance and virulence pathways for proteomic data from the polymicrobial community. A) *P. aeruginosa* (PA). B) *S. aureus* (SA). C) *S. sanguinis* (SS). KEGG pathway analysis for *P. melaninogenica* could not be performed due to a lack of available pathway data. Count is the number of proteins identified in each pathway. FDR, False Discovery Rate. Ratio is the number of identified proteins divided by the total number of proteins in the KEGG pathway.

### bEVs decrease ETI-stimulated delF508 CFTR Cl^-^ secretion, exacerbating the CF phenotype, but are not cytotoxic to pHBEC

Previously, we demonstrated that *P. aeruginosa* and bEVs secreted by *P. aeruginosa* inhibit ETI-stimulated CFTR Cl^-^ secretion by CF pHBEC (53,54). To test whether polymicrobial bEVs also inhibit CFTR Cl^-^ secretion, CFTR Cl^-^ currents were measured in wild type (WT), CF, and CF pHBEC treated with ETI. Compared with processed control treatment (PC), bEVs reduced CFTR Cl^-^ currents in WT and CF pHBEC + ETI by 39.8% (*P=*2.61E-03) and 57.2% (*P*=3.06E-05), respectively (**Fig. 5**). bEVs were non-cytotoxic to pHBEC as determined by analysis of LDH (**Fig. S1**).

**Fig. 5.**
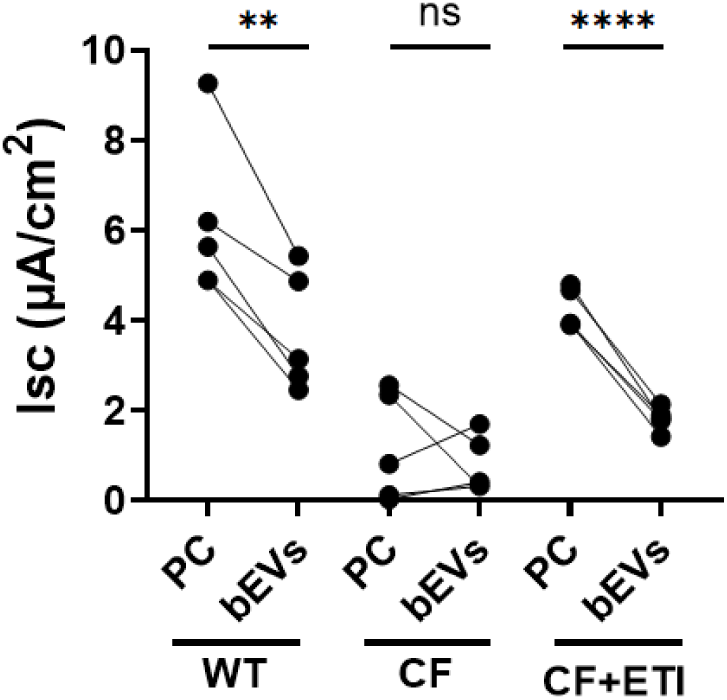
Polymicrobial bEVs decrease CFTR Cl^-^ secretion in WT and CF pHBEC treated with ETI. CFTR Cl^-^ secretion was assessed by measuring short circuit current (Isc) using Ussing chambers. WT, CF, and CF pHBEC + ETI were treated with either processed control (PC) or polymicrobial bEVs (bEVs) for 6 hrs. Significance is determined by mixed-effect linear model, with donor as a random effect. *N* = 5 donors/group. \*\**P* = 2.61E-03, \*\*\*\**P* = 3.06E-05

### Polymicrobial bEVs induce a hyperinflammatory response in CF pHBEC, which is not reverted to WT levels by ETI

To characterize the acute cellular response to bEVs, pHBEC were exposed to either PC or polymicrobial bEVs for 6 hrs., after which total RNA and protein were isolated for bulk RNA-sequencing (RNA-seq) and quantitative LC-MS/MS, respectively. Basolateral supernatants were also collected for multiplex cytokine analysis. Principal component analysis (PCA) of the RNA-seq dataset, which comprised 13,712 detected genes, revealed that samples did not cluster by bEV exposure (**Fig. 6A**). This pattern is consistent with prior reports indicating that donor-specific variation dominates over treatment effects in pHBEC (14,21,55,56). One WT pHBEC donor had a very low number of RNA read counts and was therefore excluded as an outlier. Analysis identified 163 (WT pHBEC), 114 (CF pHBEC), and 94 (CF pHBEC + ETI) differentially expressed genes (DEGs) that were significantly altered (|Log_2_(fold change)| ≥ 1, *P* < 0.05) by bEV exposure compared to PC (**Fig. 6B–D**). Notably, the three groups displayed largely non-overlapping DEGs, and no single gene was altered by bEVs in all three groups (**Fig 6E**). KEGG pathway-enrichment analysis demonstrated that ETI treatment did not revert the CF pHBEC transcriptional profile to a WT-like state (**Fig 7**). Inflammatory and metabolic pathways, including MAPK signaling, cytokine-cytokine receptor interaction, and oxidative phosphorylation, remained up-regulated in CF pHBEC + ETI (**Fig 7**), reflecting the persistent proinflammatory state and oxidative stress that characterize CF lung disease (57,58). Some KEGG pathways were differentially expressed among WT, CF, and CF pHBEC + ETI (**Fig. 7**). The MAPK pathway was increased in CF versus WT pHBEC, but unchanged by ETI treatment in CF pHBEC.

**Fig. 6.**
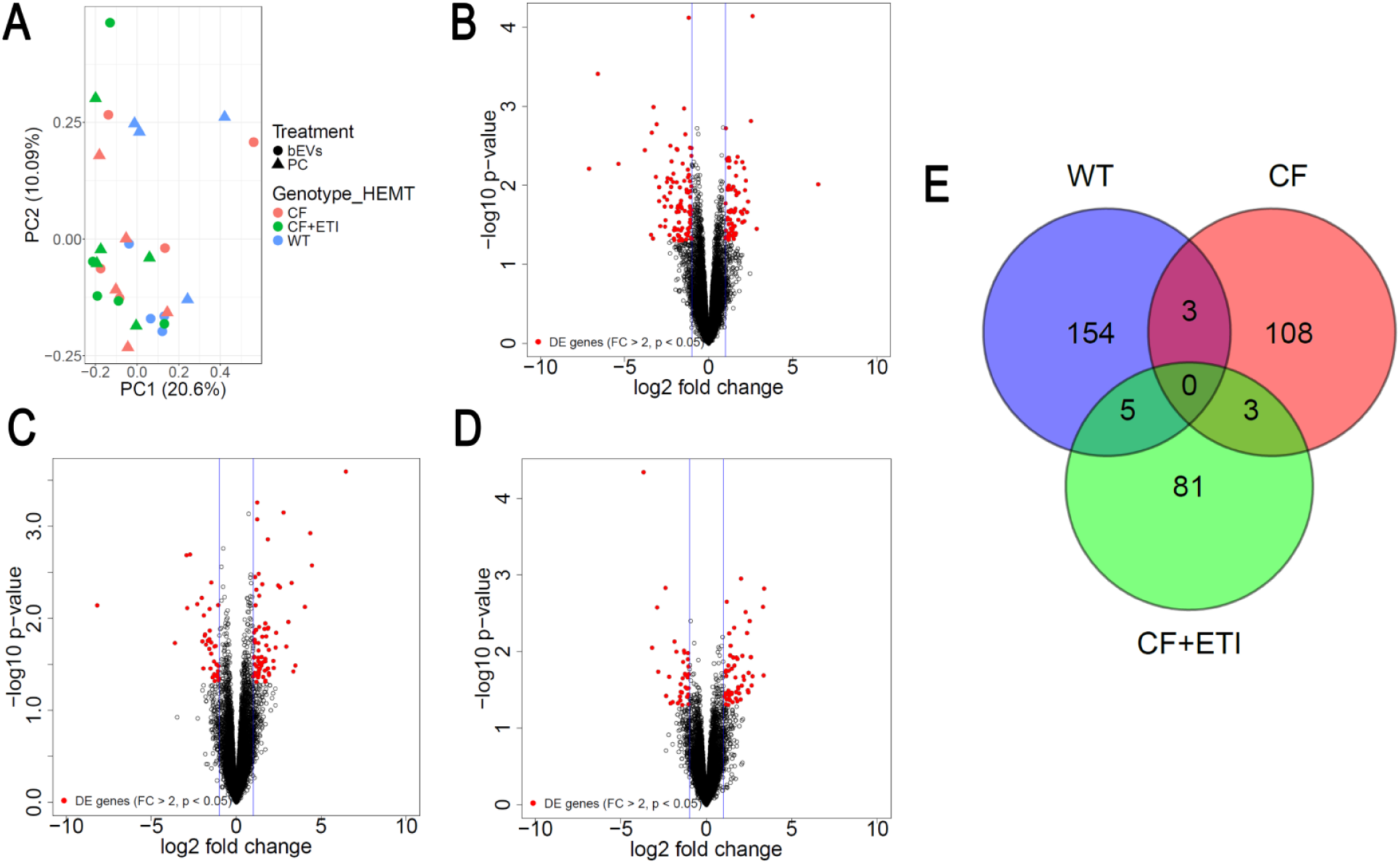
Polymicrobial bEVs induce differential gene expression in WT, CF, and CF pHBEC + ETI. Differentially expressed genes in pHBEC with bEV exposure compared to PC. Significance determined by gene-wise negative binomial generalized linear models and defined as |Log_2_(fold change)| ≥ 1 and *P* < 0.05. *N* = 5 donors/group. A) PCA plot of samples. B) WT pHBEC, full list in **Table S3**. C) CF pHBEC, full list in **Table S4**. D) CF pHBEC + ETI, full list in **Table S5**. E) Venn diagram comparing differentially expressed genes by genotype/HEMT.

**Fig. 7.**
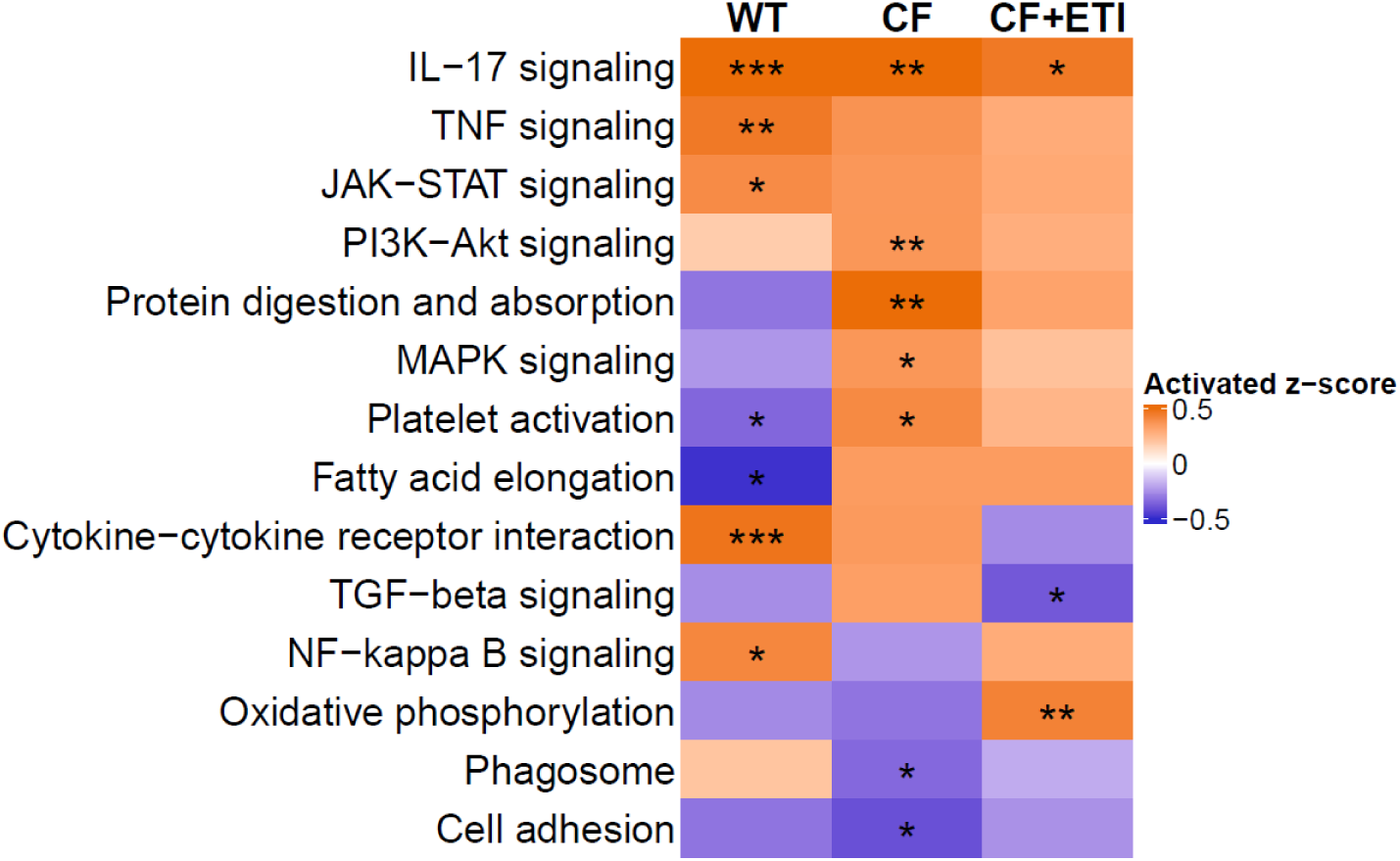
Polymicrobial bEVs induce differential pathway activation in WT, CF, and CF pHBEC +ETI. RNA-seq analysis of differentially expressed KEGG pathways in WT, CF, and CF±ETI pHBEC after 6 hrs. of bEV exposure compared to PC. Significance was determined by a mixed-effect linear model, with donor as a random effect. *N* = 5 donors/group. \**P* < 0.05, \*\**P* < 0.01, \*\*\**P* < 0.001. Exact *P* values in **Table S6**.

Proteomic profiling of pHBEC by LC-MS/MS revealed that few proteins were differentially expressed after 6 hrs. of exposure to bEVs versus PC. This finding was not unexpected since the half-life of many eukaryotic proteins is ∼9 to >20 hrs. (59,60). In WT pHBEC, only three proteins were differentially expressed after bEV exposure: ARHGAP32 (log₂FC=1.30, *P*=0.0012), a Rho-GTPase-activating protein; ADH1C (log₂FC=1.73, *P*=0.0031), an alcohol dehydrogenase; and ANKRD22 (log₂FC=-1.38, *P*=0.0032), an ankyrin-repeat domain protein (**Fig. 8A**). ANKRD22 functions as a mitochondrial Ca²⁺ regulator, and its down-regulation has been linked to excessive inflammation (61). CF pHBEC exhibited nine significantly altered proteins, including SLC4A11 (log₂FC=-1.05, *P*=0.021), which regulates oxidative-stress defenses (62) (**Fig. 8B**). The CF + ETI cohort had the most extensive proteomic response to treatment, with 73 proteins changed compared to PC (**Fig. 8C**). Among the top up-regulated proteins were RELL1 (log₂FC=1.55, *P*=1.75E-13), which induces activation of the MAPK14/p38 cascade and apoptosis when over-expressed (63), and PARD3 (log₂FC=2.04, *P*=1.75E-13), a core component of the PARD6-PARD3 complex that drives epithelial tight-junction formation (64). As with the transcriptome, there was little overlap among groups, and no protein was commonly regulated across all three pHBEC experimental groups, underscoring the genotype-and HEMT-specific nature of the bEV-induced proteomic response (**Fig. 8**).

**Fig. 8.**
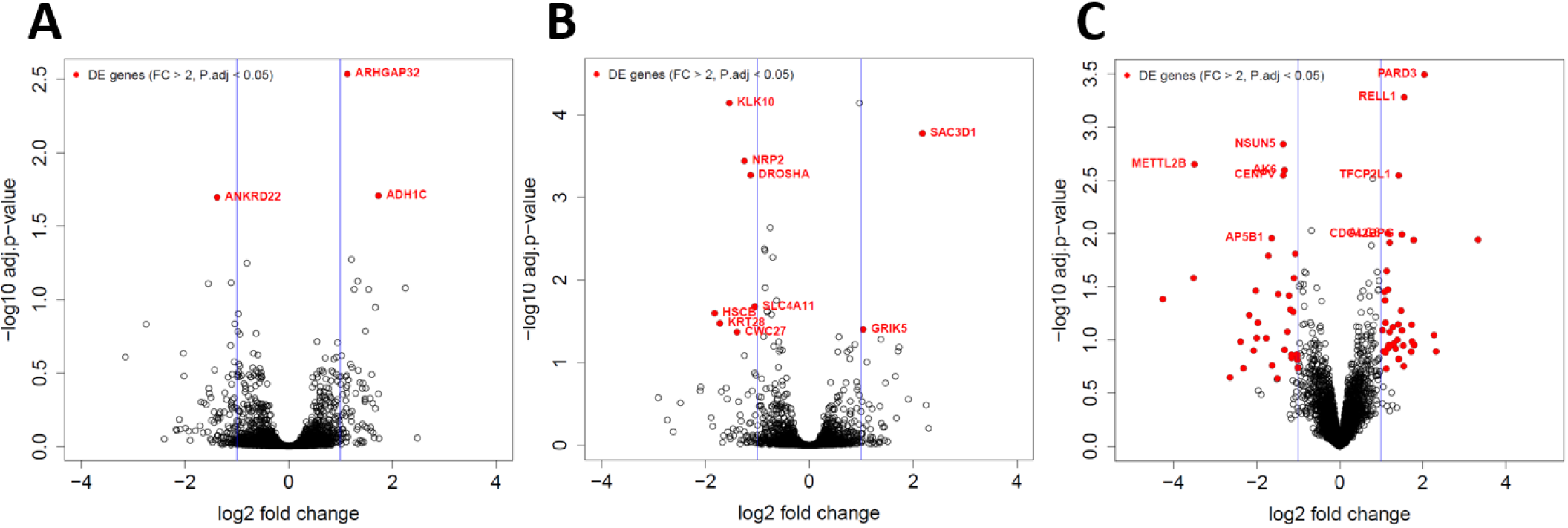
Polymicrobial bEVs induce differential protein expression in WT, CF, and CF pHBEC +ETI. Differentially expressed proteins in pHBEC with bEV exposure compared to PC. Significance determined by empirical Bayes moderated t-statistics and defined as |Log_2_(fold change)| ≥ 1 and *P* < 0.05. *N* = 3 donors/group. A) WT pHBEC. B) CF pHBEC. C) CF pHBEC + ETI.

Cytokine analysis of the basolateral media of pHBEC was consistent with the transcriptomic findings of a pro-inflammatory environment induced by bEVs. Inflammatory cytokines were increased in CF pHBEC as compared to WT pHBEC (**Fig. 9**). ETI treatment did not restore cytokines to WT levels, particularly GM-CSF, TNF-α, TGF-β, MCP-1, and IP-10 (**Fig. 9**).

**Fig. 9.**
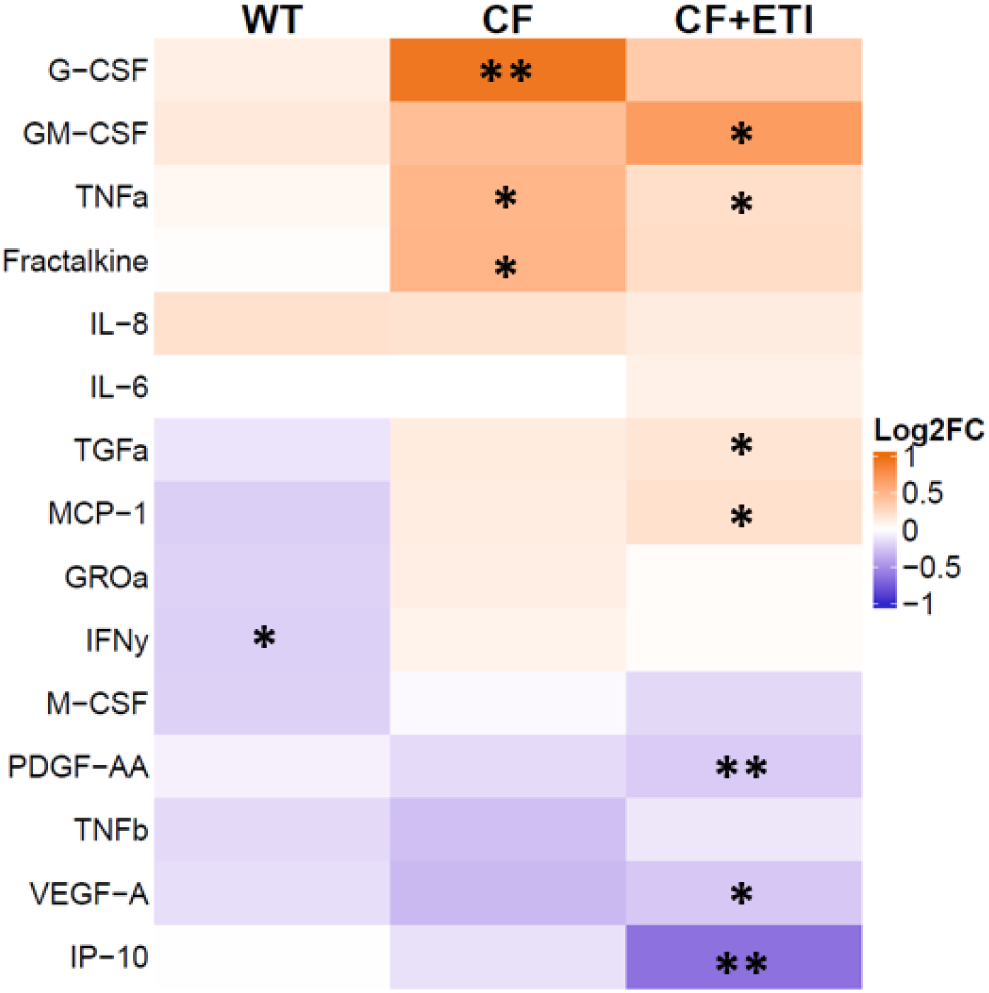
Polymicrobial bEVs elicit proinflammatory cytokine secretion by CF pHBEC. Log_2_ (Fold Change) of cytokine secretion by WT, CF, and CF pHBEC + ETI after 6 hrs. of bEV exposure compared to PC. Significance was determined by a mixed-effect linear model, with donor as a random effect. *N* = 5 donors/group. \**P* < 0.05, ** *P* < 0.01. Exact *P* values in **Table S7**.

### Polymicrobial bEVs contain sRNAs and tRNAs that inhibit host innate immunity

Previously, we demonstrated that sRNA and tRNA fragments present in bEVs secreted by *Pseudomonas* are delivered to pHBEC and are predicted to target and inhibit host mRNAs (39,43). Here, we employed a similar workflow to identify sRNAs and tRNAs in the polymicrobial bEVs. We identified a total of 67 sRNA/tRNA species by small RNA-seq that align to *P. aeruginosa,* 52 to *S. aureus*, 68 to *S. sanguinis*, and 58 to *P. melaninogenica* (**Table S8**). The 10 most abundant sRNA/tRNAs were chosen for further target prediction analysis and assessed for stable secondary structure using RNAfold (65) (**Table 1**).

**Table 1.**
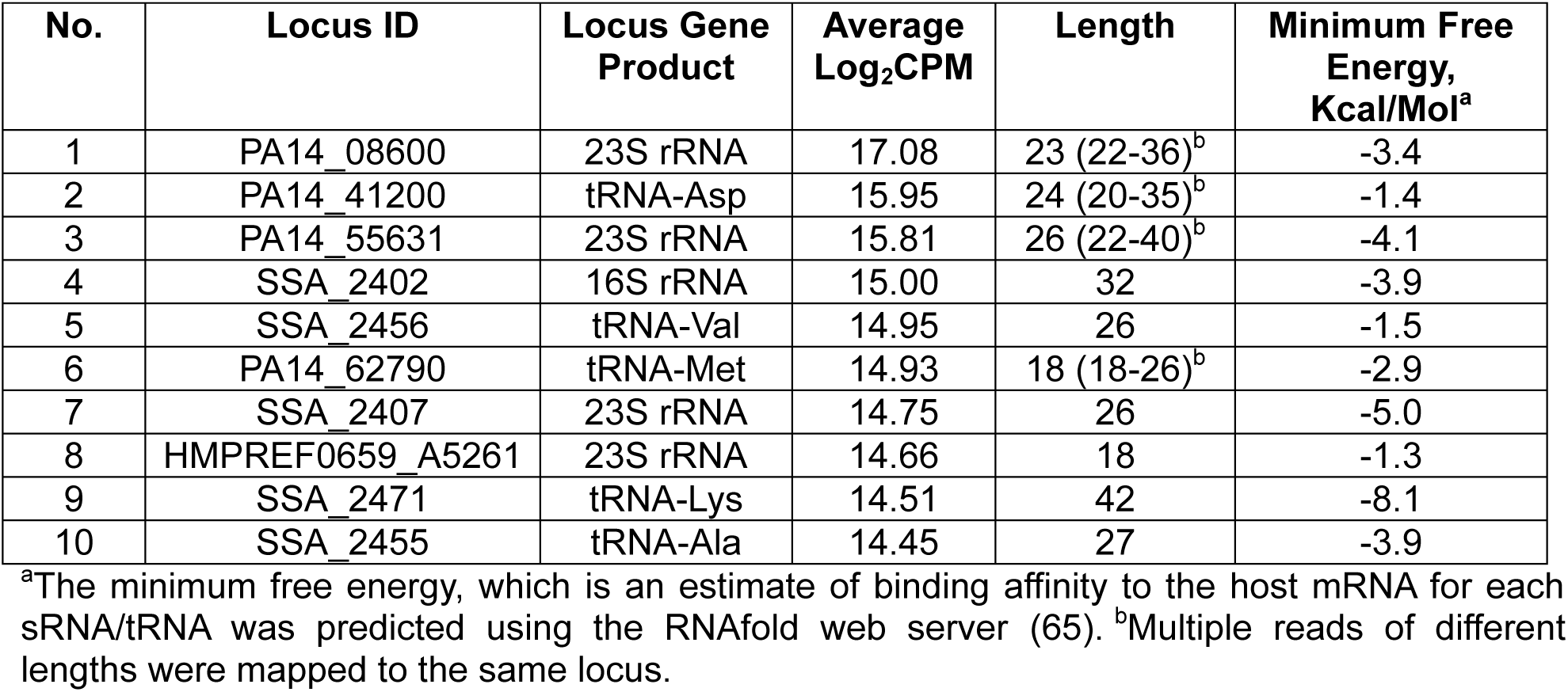
Top 10 most abundant sRNAs/tRNA fragments in polymicrobial bEVs.

| No. | Locus ID | Locus Gene Product | Average Log<sub>2</sub>CPM | Length | Minimum Free Energy, Kcal/Mol<sup>a</sup> |
| --- | --- | --- | --- | --- | --- |
| 1 | PA14_08600 | 23S rRNA | 17.08 | 23 (22-36)<sup>b</sup> | -3.4 |
| 2 | PA14_41200 | tRNA-Asp | 15.95 | 24 (20-35)<sup>b</sup> | -1.4 |
| 3 | PA14_55631 | 23S rRNA | 15.81 | 26 (22-40)<sup>b</sup> | -4.1 |
| 4 | SSA_2402 | 16S rRNA | 15.00 | 32 | -3.9 |
| 5 | SSA_2456 | tRNA-Val | 14.95 | 26 | -1.5 |
| 6 | PA14_62790 | tRNA-Met | 14.93 | 18 (18-26)<sup>b</sup> | -2.9 |
| 7 | SSA_2407 | 23S rRNA | 14.75 | 26 | -5.0 |
| 8 | HMPREF0659_A5261 | 23S rRNA | 14.66 | 18 | -1.3 |
| 9 | SSA_2471 | tRNA-Lys | 14.51 | 42 | -8.1 |
| 10 | SSA_2455 | tRNA-Ala | 14.45 | 27 | -3.9 |

<sup>a</sup>The minimum free energy, which is an estimate of binding affinity to the host mRNA for each sRNA/tRNA was predicted using the RNAfold web server (65). <sup>b</sup>Multiple reads of different lengths were mapped to the same locus.

Potential gene targets in WT, CF, and CF + ETI pHBEC were predicted for each candidate sRNA and tRNA using the miRanda algorithm (**Table 2**). On average, the most abundant sRNAs/tRNAs are predicted to regulate 17.6% of DEGs in WT pHBEC, 15% in CF pHBEC, and 17.1% in CF + ETI pHBEC (**Table 2**). These 10 most abundant sRNAs and tRNA fragments reveal a potential mechanism of transcriptional regulation of pHBEC by polymicrobial bEVs. Notably, tRNA PA14_62790, which reflects a stress response in *P. aeruginosa* (39) was abundant in bEVs (**Table 1**). We have previously shown *in vivo* that this tRNA downregulates IL-8 and neutrophilic airway inflammation in response to infection to decrease the clearance of bacteria (39).

**Table 2.**
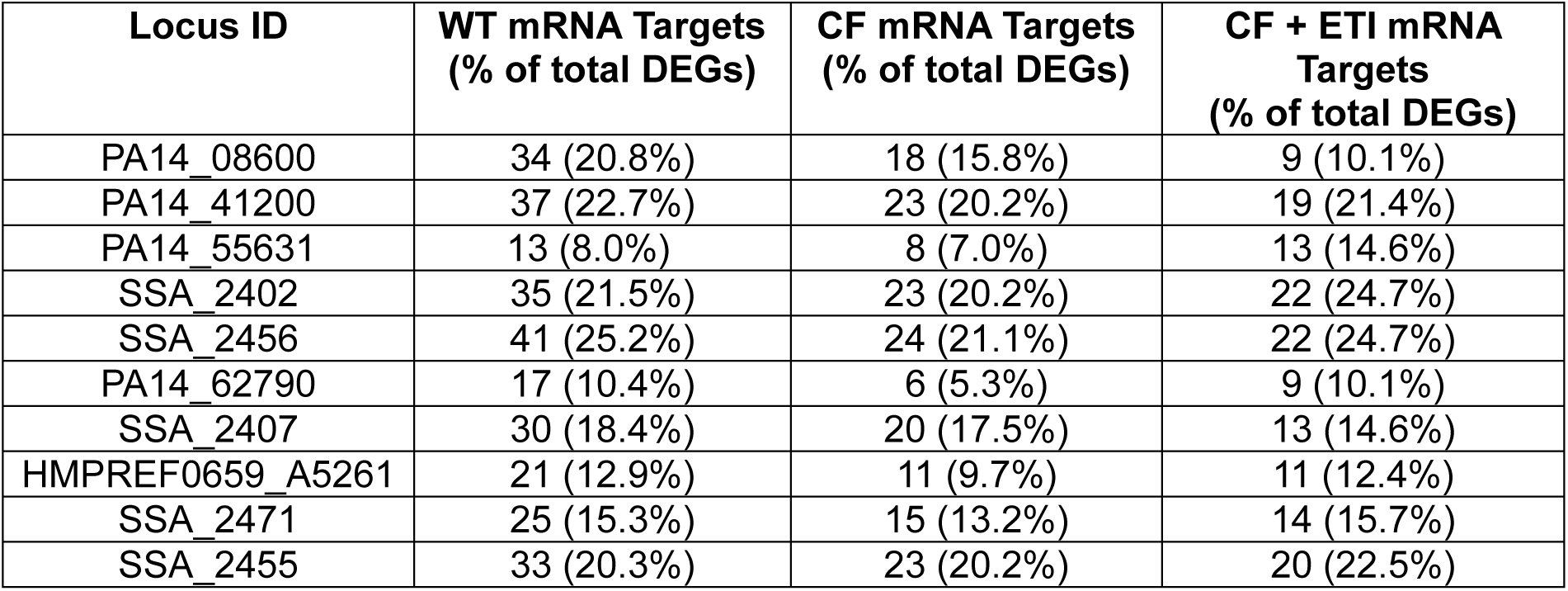
Number of differentially expressed mRNAs in pHBEC that are predicted targets of sRNA/tRNA fragments.

| Locus ID | WT mRNA Targets<br>(% of total DEGs) | CF mRNA Targets<br>(% of total DEGs) | CF + ETI mRNA<br>Targets<br>(% of total DEGs) |
| --- | --- | --- | --- |
| PA14_08600 | 34 (20.8%) | 18 (15.8%) | 9 (10.1%) |
| PA14_41200 | 37 (22.7%) | 23 (20.2%) | 19 (21.4%) |
| PA14_55631 | 13 (8.0%) | 8 (7.0%) | 13 (14.6%) |
| SSA_2402 | 35 (21.5%) | 23 (20.2%) | 22 (24.7%) |
| SSA_2456 | 41 (25.2%) | 24 (21.1%) | 22 (24.7%) |
| PA14_62790 | 17 (10.4%) | 6 (5.3%) | 9 (10.1%) |
| SSA_2407 | 30 (18.4%) | 20 (17.5%) | 13 (14.6%) |
| HMPREF0659_A5261 | 21 (12.9%) | 11 (9.7%) | 11 (12.4%) |
| SSA_2471 | 25 (15.3%) | 15 (13.2%) | 14 (15.7%) |
| SSA_2455 | 33 (20.3%) | 23 (20.2%) | 20 (22.5%) |

Together, our analyses demonstrate that polymicrobial bEVs elicit distinct transcriptional, proteomic, and secretory responses in pHBEC, and that ETI exposure of CF pHBEC does not fully mitigate the bEV-driven activation of inflammatory and stress-response pathways that perpetuate CF lung damage and the establishment of chronic infection.

## DISCUSSION

In this study, we report a new host-pathogen infection model developed to elucidate the effects of polymicrobial bEVs on pHBEC and to assess the effects of ETI on CF pHBEC under physiological conditions. *P. aeruginosa*, *S. aureus*, *S. sanguinis*, and *P. melaninogenica* were selected based on our previous study using K-means clustering, the gap statistics machine learning approach, and metabolic modeling of 16S rRNA data from a cohort of 167 people with mild to moderate CF (23). The polymicrobial genera were identified in 34% of pwCF and account for 27% of the variability in lung function in our cohort (23). Variables, such as a single bacterial genus, Simpson diversity, and age, explain less than 11% of the variability in patient lung function (23). Moreover, the elimination of one or more bacterial genera from our model dramatically reduces the number of pwCF in our cohort with those infections, whereas the addition of genera does not increase the number of pwCF with those infections (23). The polymicrobial culture is grown in artificial CF sputum under anoxic conditions designed to resemble anoxic mucus plugs overlying pHBEC. As bacteria reside primarily in the mucus (35), communication between bacteria and host cells, in part, is mediated by the secretion of bEVs (39,43,53,66–69). Our 16S rRNA characterization of bEVs isolated from CF BALF obtained at Dartmouth Health identified bEVs secreted by *Pseudomonas*, *Staphylococcus*, *Streptococcus*, and *Prevotella*, which represented approximately 53% of the entire microbial community detected in BALF samples. Thus, we developed a biologically relevant polymicrobial infection model to study the interactions between bEVs and pHBEC.

Using a variety of approaches, including RNA-seq and proteomics of bacteria, bEVs, and pHBEC, cytokine secretion, and CFTR Cl^-^ currents by pHBEC, we have made several novel observations: (1) bEVs contain proteins, sRNAs, and tRNA fragments that are predicted to increase infection and stimulate pro-inflammatory pathways in CF pHBEC; (2) the pathogenic effects of bEVs, include reducing ETI-stimulated *Phe508del/Phe508del* CFTR Cl^-^ currents, and (3) ETI failed to reduce the hypersecretion of pro-inflammatory cytokines when exposed to bEVs. These observations extend our earlier work on *P. aeruginosa* bEVs, in which we identified protein, sRNA, and tRNA fragments that suppress the host immune response to favor infection and exacerbate the CF phenotype (39,43,54)

To characterize the polymicrobial bEVs, we applied orthogonal methods, including 16S rRNA sequencing, fluorescence-based quantification, RNA-seq, and proteomics. Although the lack of available species-specific antibodies limits direct measures of bEV abundance, the consensus between these independent approaches provides confidence that all four bacteria secrete bEVs. We acknowledge that RNA cargo may be packaged differently across subpopulations of bEVs, potentially biasing any single quantification method. However, the general agreement across methods mitigates this concern and allows us to proceed with functional analyses of bEV content. bEV cargo was enriched for proteins implicated in biofilm formation, quorum sensing, and β-lactam resistance, processes that facilitate chronic infection in the CF lung (**Fig. 3**). bEVs also contain proteins, sRNA, and tRNA fragments that, as shown in previous studies, are delivered to pHBEC and are predicted to target genes and numerous KEGG pathways, including NFκB (39,43,66,67).

Polymicrobial bEVs eliminated the ability of ETI to stimulate CFTR Cl^-^ currents in CF pHBEC (**Fig. 5**), which would reduce the efficacy of ETI (53). Previously, we showed that *P. aeruginosa* bEVs decrease CFTR Cl⁻ secretion, exacerbating the CF phenotype (53). By contrast, *S. aureus* and *Streptococcus* spp. alone have no effect on CFTR Cl^-^ secretion (70). The effect of bEVs released by *S. sanguinis* and *P. melaninogenica* in monoculture on CFTR Cl^-^ currents remains unknown, and additional studies are needed to address this gap. Ideally, one would isolate bEVs from each organism within the polymicrobial community to assess their specific effects, but this lies beyond the scope of the present study. While it is feasible to analyze bEVs derived from monocultures, their protein and sRNA/tRNA fragment cargo is likely to differ from that of bEVs produced in a mixed community, as inter-bacterial interactions in coculture markedly influence gene expression and bacterial phenotype (22,25–27). Moreover, *Prevotella* fails to grow in SCFM2 unless co-cultured with *Pseudomonas* or *Staphylococcus* (22,27), making it impossible to obtain *Prevotella*-derived bEVs from monocultures in this medium.

To further determine how bEVs affect epithelial physiology, we examined transcriptional and proteomic responses of pHBEC to bEVs. Exposure to polymicrobial bEVs triggered genotype and ETI-specific changes in gene expression, with a pronounced upregulation of pro-inflammatory pathways (**Fig. 7**). These findings reinforce the need for additional targeted strategies to dampen lung inflammation in pwCF. Supporting this, a recent publication has shown that lung inflammation persists in pwCF treated with ETI after 1.5 years of treatment (71). As an initial step toward pinpointing therapeutic targets, we mapped polymicrobial bEV-associated sRNA/tRNA fragments to host response genes differentially regulated by bEVs. We identified sRNAs and tRNAs predicted to regulate approximately 17% of DEGs during bEV exposure (**Table 2**), highlighting potential therapeutic targets for future study.

Our study has a few limitations. First, it would be advantageous to study the effects of the polymicrobial culture in an animal model of CF and on pHBEC in addition to bEVs. Bacteria secrete additional factors absent in bEVs that affect the host and may also alter their transcriptional profile and bEV secretion when co-incubated with pHBEC, a trait not captured in our reductionist bEV treatment model. Second, most of the bEV sRNAs we predict to interact with pHBEC remain unvalidated experimentally. Follow-up experiments will be required to confirm functional relevance. However, we argue that computational predictions are a powerful tool to guide *in vitro*/*in vivo* experimentation (39). Our study has several novel advantages including: (1) We use a newly-developed CF polymicrobial community to characterize bEVs; (2) bEVs were characterized using orthogonal approaches and the TEM experiments were conducted by an investigator blinded to sample identity; (3) Primary HBEC cells from 5 WT and 5 CF donors were studied, which captures some of the variability between pwCF, as compared to a cell line obtained from a single donor, thereby enhancing experimental rigor; and (4) We conducted RNA-seq and proteomics on bEVs and pHBEC as well as measurements of cytokine and CFTR Cl^-^ secretion by pHBEC.

In conclusion, we have shown that our polymicrobial community secretes bEVs that contain bacterial signaling molecules predicted to promote biofilm formation and antibiotic resistance within the polymicrobial community, amplify the secretion of pro-inflammatory cytokines, and inhibit ETI-stimulated CFTR Cl^-^ secretion in CF pHBEC. Interestingly, these negative effects of bEVs were not fully mitigated by ETI, underscoring the need for (1) additional studies characterizing the impact of polymicrobial bEVs on the pathogenesis of CF lung infections and (2) the development of new therapies to reduce host inflammation and inhibit CFTR-suppressive pathways by CF pHBEC. We anticipate that this study will facilitate the development of more effective approaches to eliminate infection and inflammation in CF and other lung diseases characterized by chronic polymicrobial infections and excessive inflammation.

## MATERIALS AND METHODS

### Polymicrobial culture

The polymicrobial culture containing *Pseudomonas aeruginosa* PA14, *Staphylococcus aureus Newman*, *Streptococcus sanguinis* SK36, and *Prevotella melaninogenica* ATCC 25845 was cultured in Synthetic Cystic Fibrosis Medium 2 (SCFM2)(72–74) under 0% O_2_, as described previously (22,74). After incubation in 0% O_2_ for 24 hrs. at 37°C, cultures were collected for bEV isolation as described below. Aliquots of the polymicrobial cultures were plated onto selective media to count colony-forming units (CFUs).

### Isolation and characterization of bEVs

bEVs were isolated as described previously using OptiPrep density gradient ultracentrifugation (39,40,43,53,54,68). 500 µL fractions were removed starting from the top of the resulting density gradient, and average particle size and concentration were measured with a Nanosight NS300 using a 532 nm laser, camera level 15, detection threshold 7, variable focus, and screen gain 10. Three thirty-second videos per sample were captured for analysis. Protein concentration of the bEVs, to assess bEV purity (45), was measured using a Pierce BCA™ Protein Assay Kit (Thermo Fisher Scientific, Cat. No. 23225).

### Transmission Electron Microscopy of bEVs

Negative-staining TEM was used to visualize PC and polymicrobial bEV preparations, as recommended by the ISEV (44), and previously described in detail (39). Fractions 1 and 2 from the PC or bEV OptiPrep gradients were combined and concentrated for 2 hrs. by ultracentrifugation at 39,000 g at 4°C before sample preparation. Microscopy images were processed in MetaMorph Offline image analysis software (v.7.8.0.0; Molecular Devices LLC, San Jose, CA), where the diameters of bEVs were measured by drawing a line across each vesicle. All imaging was done by an investigator blinded to the sample identity.

### Characterization of bEVs by 16S sequencing

DNA from bEVs was isolated using the QIAPrep Spin Mini Prep 250 kit (Qiagen, Germantown, MD, Cat. No. 27106) with lysozyme (800 µg/mL) and lysostaphin (12.5 µg/mL) to lyse *S. aureus* vesicles. DNA concentrations were determined by Qubit fluorometry prior to library preparation and 16S sequencing (SEQCENTER LLC, Pittsburgh, PA). Samples were prepared using Zymo Research’s Quick-16S kit with phased primers targeting the V3/V4 regions of the 16S gene. After cleanup and normalization, samples were sequenced on a P1 600cyc NextSeq2000 Flowcell to generate 2×301bp paired-end (PE) reads. Quality control and adapter trimming were performed with bcl-convert (v4.2.4). Sequences were imported to Qiime2 (v2023.5.1 (q2cli) from quay.io/qiime2 Docker Image) for analysis, and primer sequences were removed using Cutadapt2. Sequences were denoised using dada2. Denoised sequences were assigned operational taxonomic units (OTUs) using the Silva 138 99% OTUs full-length sequence database and the VSEARCH4 utility within Qiime2’s feature-classifier plugin. OTUs were then collapsed to their lowest taxonomic units, and their counts were converted to reflect their relative frequency within a sample.

### Characterization of bEVs by fluorescence

To identify bacterial bEVs using an orthogonal approach to 16S sequencing, bEVs secreted by *P. aeruginosa-*CFP, and *S. aureus*-GFP were isolated as described above. *S. sanguinis* was stained with CellTracker Deep Red fluorescent probe (Thermo Fisher Scientific, Cat. No. C34565), and *P. melaninogenica* was stained with CellTracker Orange CMTMR (Thermo Fisher Scientific, Cat. No. C2927) according to the manufacturer’s protocol before isolation of bEVs as described above. Particle counting and fluorescence of the polymicrobial bEVs were measured on a ZetaView PMX-430-Z QUATT system 405/488/520/640 (Particle Metrix, Meerbusch, Germany).

### Primary Human Bronchial Epithelial Cells (pHBEC)

pHBEC from nonsmoker donors aged 14-54 years were obtained from five WT (3 males and 2 females) and five CF (*Phe508del/Phe508del*, 1 male and 4 females), and were cultured and polarized on filters at air-liquid interface according to published protocols (53,75). For CF pHBEC, ETI (Elexacaftor/Tezacaftor/Ivacaftor; 3 µM VX-445, 3 µM VX-809, 100 nM VX-770) or DMSO (vehicle control) was added to the basolateral media 48 hrs. before experiments. The Dartmouth Committee for the Protection of Human Subjects concluded that the use of pHBEC in this study is not considered human subject research as the cells were taken from discarded tissue with informed consent and do not contain any patient-identifying information.

### Exposure of pHBEC to bEVs

pHBEC were exposed apically to either PC or polymicrobial bEVs (2×10^10^/mL per filter, concentrations of bEVs measured in sputum of pwCF (76)). pHBEC were then incubated for 6 hrs. at 37°C with 5% CO_2_ /21% O_2_/balance N_2_ before being washed twice with PBS supplemented with 1 mM MgCl_2_ and 0.1 mM CaCl_2_ (pH 8.2) and immediately processed for experiments.

### Analysis of CFTR Cl^-^ currents by pHBEC

CFTR Cl^-^ currents were measured by Ussing chambers and described in detail in previous studies (53,70,77). Data are expressed as the forskolin-stimulated, CFTR_inh_-172-inhibited short circuit current (Isc), presented as μA/cm^2^.

### Cytotoxicity of polymicrobial bEVs

The CytoTox 96® Non-Radioactive LDH Cytotoxicity Assay (Promega, Cat. No. G1780) was performed according to the manufacturer’s instructions.

### RNA isolation, bulk RNA-seq, and downstream analyses

RNA-seq and data analysis were performed as described previously (78). Reads were aligned to the GRCh38.97 human genome. The edgeR package (v4.4.2) was used for count filtering, normalization, and differential gene expression analysis (79,80). Transcripts with fewer than 10 counts per library were filtered out, retaining 13,712 genes. Sequencing data have been deposited to the National Center for Biotechnology Information Gene Expression Omnibus (NCBI GEO) (GSE325885).

### Small RNA-seq (<200 nucleotides) and downstream analyses

Libraries were prepared using the NEBNext Low-bias Small RNA Library Prep Kit (New England Bio Labs, Cat. No. E3420L) according to the manufacturer’s instructions. Libraries were pooled and sequenced at a depth of 10 million single-end reads on a NextSeq2000 platform. Reads were trimmed with Cutadapt (v4.0) (81) and aligned to the *P. aeruginosa* UCBPP-PA14 (NC_008463.1), *S. aureus* Newman (NC_009641.1), *S. sanguinis* SK36 (NC_009009.1), and *P. melaninogenica* ATCC 25845 (NC_014370.1) transcriptomes using Hisat2 (v2.2.1) (82). sRNA expression levels were quantified with featureCounts (v2.0.1) (83), using the respective Refseq annotation assemblies. Final analyses and figures were generated in R using ggplot2 (v4.0.1). RNA secondary structure predictions and minimum free energy calculations were obtained using the RNAfold WebServer (65). The miRanda microRNA target-scanning algorithm (v3.3a)(84) with a minimum alignment score of 150 was used to predict human target genes of sRNAs. Sequencing data have been deposited to NCBI GEO (GSE325886).

### Proteomic analysis of pHBEC and bacteria

Proteins were isolated from pHBEC treated with PC or bEVs for 6 hrs. Samples were prepared as described previously (85). For mass spectrometry analysis, dried peptides were resuspended in 0.1% formic acid in water (LC-MS grade, Fisher Scientific), and eptides were analyzed using nanoElute2 coupled with timsTOF Pro2 Mass Spectrometer (Bruker Daltonics). Solvent A was 0.1% FA in water (LC-MS grade, Fisher Scientific), and Solvent B was 0.1% FA in ACN (LC-MS grade, Fisher Scientific). Peptides were separated on a PepSep C18 column (25 cm length, 150 μm I.D., 1.5 μm particle size, 100 Å pore size; Bruker Daltonics), at 600 nL/min flow rate, using a linear gradient of 3% to 25% of Solvent B in 48 min, then raised to 35% B at 54 min, followed by column washing with 95% B. Total method run time was 60 minutes. The source parameters were: capillary voltage = 1700 V, dry gas = 3.0 L/min, and dry temperature = 180 °C. Both the ramp and accumulation times were set to 75 ms. The dia-PASEF method was designed using the py_diAID tool. Briefly, the data was collected in m/z range 300-1200 m/z and mobility range (1/K0) = 0.65-1.35 Vs cm−2 with the following parameters: number of MS1 ramps = 1, number of MS/MS ramps = 14, number of MS/MS windows = 28. The estimated cycle time was 1.20 s. The raw data files were analyzed by Spectronaut (v. 18.7) using default BGS factory settings in a library-free manner (directDIA+). The human SwissProt database was used for pHBEC samples, and a combined database of *P. aeruginosa* PA14, *P. aeruginosa* PA01, *S. aureus* Newman, *S. sanguinis* SK36, and *P. melaninogenica* was used for the polymicrobial bEVs and whole bacteria samples. Peptide spectrum matches, peptides, and protein groups were identified using a false discovery rate (FDR) threshold of 0.01. Proteomics data have been deposited in the ProteomeXchange Consortium via the PRIDE (86) partner repository (PXD076596).

Resulting protein intensities were analyzed for differential expression with the DEP2 package (87) in R (v4.4.0). Protein expression was filtered and normalized according to default parameters, and missing values imputed with the “MinDet” setting. Significant proteins were defined as having an absolute Log_2_(fold change) ≥ 1 and an unadjusted *P* < 0.05 and marked by add_rejections. bEV proteins were analyzed for KEGG pathway activation by ESKAPE Act Plus (50).

### Cytokine secretion by pHBEC

Immediately after the 6 hr. treatment of pHBEC with PC or bEVs, 500 µL of basolateral medium was aliquoted for cytokine analysis. As a carrier protein, 55 µL of 5% BSA was added per sample before storage at -80°C. Cytokines were measured using Millipore human cytokine multiplex kits (48-plex; EMD Millipore Corporation, Billerica, MA) according to the manufacturer’s instructions.

### Statistics

Data were analyzed for statistical significance in R (v4.4.0). Respective statistical analyses and *P* values are detailed within figure legends. Differential gene expression for RNA-seq data was determined using gene-wise negative binomial generalized linear models using edgeR. Differential protein expression was determined using the DEP2 R package. KEGG analyses in ESKAPE Act Plus utilize binomial tests to assess pathway significance, as described (50). All other analyses were conducted using mixed-effect linear models with donor as a random effect, which we have used extensively in similar studies (43,88,89).

## ACKNOWLEDGEMENTS

Research in this study was supported by the Cystic Fibrosis Foundation (STANTO19R0), the National Institutes of Health (R01HL151385, R01HL174700, and P30DK117469), and the Flatley Foundation to BAS, NIH F31HL172440 to LAC, and NIH T32AI007519. Imaging was performed at Microscopy Imaging and Cytometry at the University of Vermont (RRID:SCR_018821). RNA-seq data analysis support was provided through the Genomics and Molecular Biology Shared Resource (RRID:SCR_021293) at Dartmouth by the NCI Cancer Center (P30CA023108) and NIH (1S10OD030242), and Dartmouth’s Center for Quantitative Biology (CQB) Genomics Data Sciences Core (GDSC) supported by the NIH National Institute of General Medical Sciences (3P20GM130454). Cytokine secretion experiments were carried out in DartLab, the Immune Monitoring and Flow Cytometry Shared Resource at the Norris Cotton Cancer Center at Dartmouth, with NCI Cancer Center Support Grant 5P30 CA023108-41. Mass Spectrometry analyses were performed by the Mass Spectrometry Technology Access Center at the McDonnell Genome Institute (MTAC@MGI) at Washington University School of Medicine, supported by the Diabetes Research Center/NIH grant P30 DK020579, Institute of Clinical and Translational Sciences/NCATS CTSA award UL1 TR002345, and Siteman Cancer Center/NCI CCSG grant P30 CA091842. We thank Dr. Jennifer Bomberger for the *P. aeruginosa*-CFP, the *S. aureus*-GFP, and the protocols for staining with CellTracker fluorescent probes.

